# Ultrastructure Expansion Microscopy in *Trypanosoma brucei*

**DOI:** 10.1101/2021.04.20.440568

**Authors:** Ana Kalichava, Torsten Ochsenreiter

## Abstract

The recently developed ultrastructure expansion microscopy (U-ExM) technique allows to increase the spatial resolution within a cell or tissue for microscopic imaging through the physical expansion of the sample. In this study we validate the use of U-ExM in *Trypanosoma brucei* by visualizing the nucleus and kDNA as well as proteins of the cytoskeleton, the basal body, the mitochondrion and the ER. *T. brucei* is a unicellular flagellated protozoan parasite and the causative agent of human African sleeping sickness and Nagana in cattle.The highly polarized parasite cell body is about 25 μm in length and is shaped by the subpellicular microtubule corset. Its single flagellum emanates from the posterior part of the cell and is attached along the entire cell body. *T. brucei* The cell contains all typical organelles of eukaryotic cells including ER, Golgi and mitochondrion. Interestingly, Golgi and mitochondrion are single unit organelles in this protozoan parasite. The signature feature of trypanosomes is the single unit mitochondrial genome, the kinetoplast DNA (kDNA) that is organized in a complex structure of interlocked mini- and maxicircles. The kDNA is segregated during cell division by the tripartite attachment complex (TAC) that connects it via the mitochondrial membranes to the base of the flagellum.

## Results

### 1. Ultrastructure expansion microscopy, cell isotropicity and expansion factor in *T. brucei*

Expansion microscopy is based on a series of chemical steps to physically magnify the cell and visualize ultrastructures by optical microscopy (for details see [1–3]). In the first step the precursor molecules formaldehyde and acrylamide are introduced to functionalize the cellular components allowing the subsequent linkage of swellable polyelectrolyte gel (sodium acrylate). The gel formation is accomplished analogously to polyacrylamide gels using oxidizing reagents like tetramethylethylenediamine and ammonium persulfate. After the gel is formed, the cell is denatured using sodium dodecyl sulfate and heat and is then ready for the application of standard immunofluorescence techniques. These approaches include antibody based localization of proteins and DNA stains such as DAPI. Most importantly the gel embedded cells can now be expanded (Figure 1A). For *T. brucei* we are now routinely achieving expansion with a factor of 4.5 (Figure 1B). The use of ultrastructure expansion microscopy was first demonstrated in Kinetoplastea localizing a component of the mitochondrial genome segregation machinery (TAP110) relative to its interaction partner TAC102 [4]. In that study the isotropicity was measured using the basal body, the kDNA and the nucleus as references. The expansion factors of these cellular compartments/complexes varied less than 10% between 3.61 and 3.86. In the current study we were able to increase the expansion factor to 4.2 - 4.6 as measured by the expansion of the kDNA, nucleus and cytoskeleton (Figure 1C-D). Aside from the measurements of cellular compartments the isotropicity of the expansion becomes obvious in the maintenance of the overall cell shape when compared to the non expanded trypanosomes (Figure 1B).

**Figure 1:**
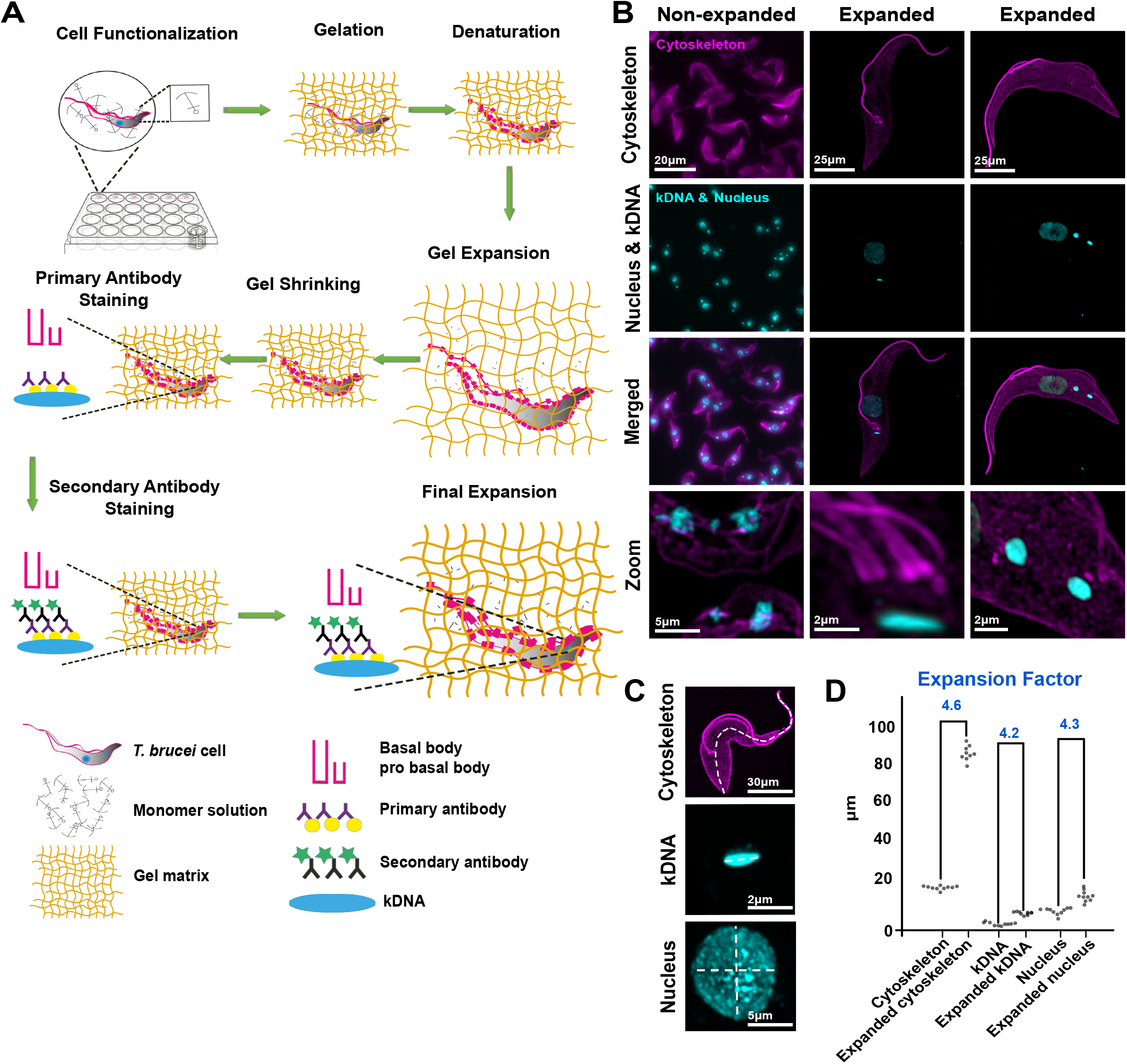
Schematic representation of ultrastructure expansion microscopy concept and example images. (**A**) Diagram representing the workflow for U-ExM in *T. brucei*. First, cells were settled at room temperature on coverslips. Then, the coverslips were incubated in 24 well-plate filled with a solution of formaldehyde and acrylamide in phosphate-buffer saline (PBS). Cells were subsequently prepared for gelation by carefully transferring coverslips into a gel solution supplemented with ammonium persulfate (APS) and Tetramethylethylenediamine (TEMED). After gelation, the cell components were denatured using heat and Sodium Dodecyl Sulfate (SDS), which was followed by first round expansion and gel shrinkage in PBS. Then, cells were stained with primary and secondary antibodies followed by dializing in water for the final expansion. (**B**) Immunofluorescence confocal microscopy imagery of non-expanded and expanded *T. brucei* cells (α-tubulin, magenta); kDNA and nuclear DNA (cyan). Imagery was deconvolved using Huygens Professional and visualized using Imaris. (**C-D**) Measurements of cytoskeleton length, kDNA length and nucleus diameter in non-expanded and expanded cells were done in ImageJ. Expansion factor was calculated as the ratio between non-expanded and expanded cell compartments.

### 2. The kDNA

The ability to image DNA in the nucleus and the mitochondrion is of general interest and thus we evaluated the use of the minor groove AT-rich DNA binder DAPI in U-ExM. The mitochondrial DNA of most Kinteoplastea is organized in a complex structure of interlocked minicircles and maxicircles resulting in the DAPI stain to appear as a homogeneous rod/disc like structure [5]. As a consequence of minicircles being covalently interlocked in U-ExM the kDNA disc diameter is isotropically expanded but it remains largely unchanged in its height. Using U-ExM we can visualize individual regions within the kDNA disc that have previously only been detected by electron microscopy. This includes the inner unilateral filament region, a DNA containing domain between the kDNA disc and the inner mitochondrial membrane (Figure 2A) [6].

**Figure 2:**
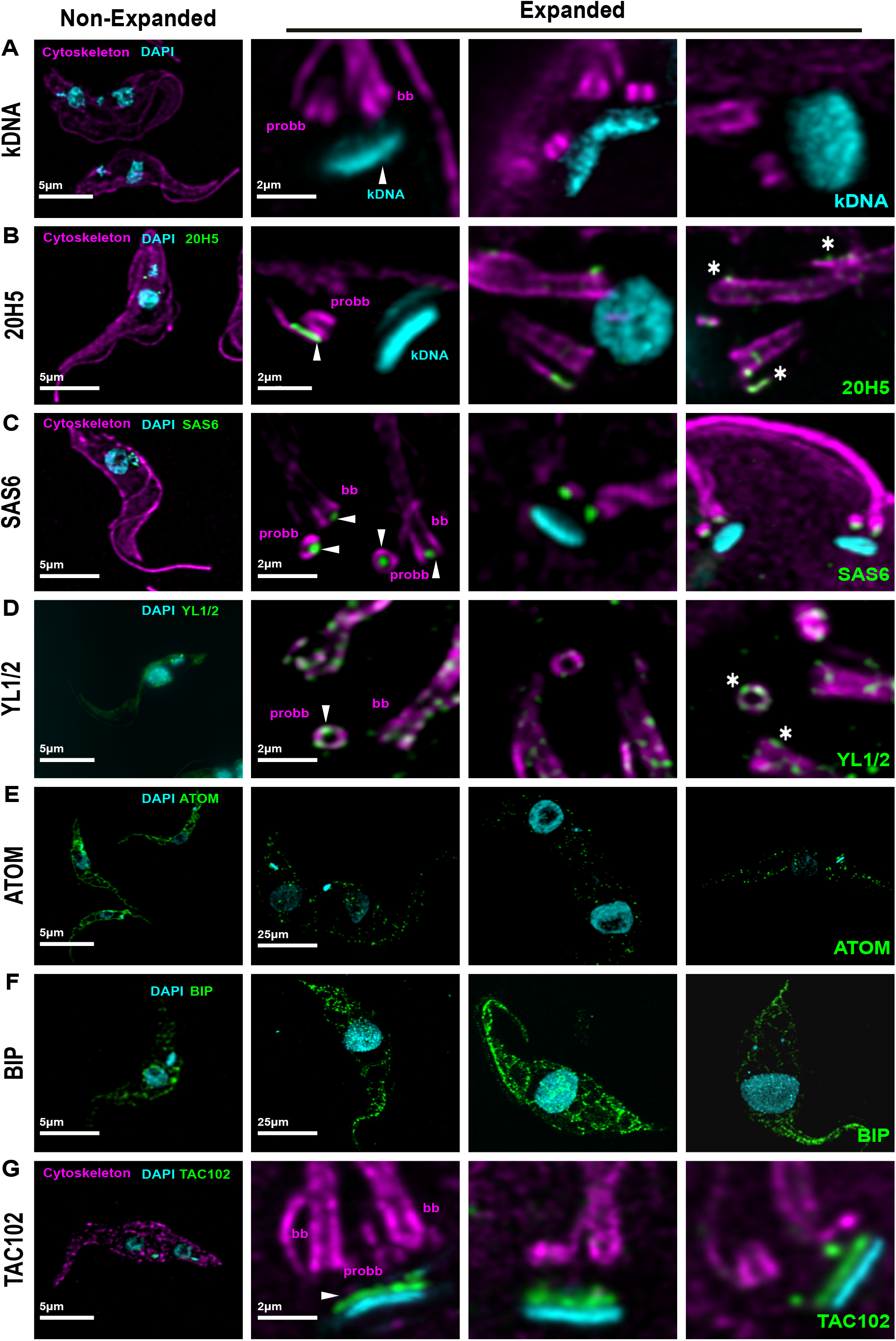
Nanoscale resolution imaging of different compartments of *T. brucei*. Representative comparative confocal images of expanded and non-expanded *T. brucei*. (**A**) kDNA (DAPI, cyan) images during different stages (segregation, replication and top view) of the cell cycle. (**B**) Centrin stained with anti centrin antibody 20H5 (green), anti α-tubulin antibody (magenta), kDNA and nucleus with DAPI (cyan). (**C**) Nine homodimer cartwheel protein anti SAS6 antibody (green) localisation at the bb and probb (magenta). (**D**) Tyrosinated tubulin distribution (YL1/2, green) at the bb and probb (magenda). (**E**) Mitochondrial protein complexes (anti ATOM antibody, green) distribution in *T. brucei*. (**F**) BIP (anti BIP antibody, green) distribution in *T. brucei*. (**G**) TAC protein (anti-TAC102 monoclonal antibody, green) localization at the kDNA. All microscopic images were deconvoluted in Huygens Professional and analyzed in Imaris.

### 3. The cytoskeleton and the basal body

In *T. brucei* the cytoskeleton is dominated by microtubules. We visualized the cell body using a commercial alpha-tubulin antibody (Sigma Adlrich, 04-1624). With this reagent we can follow the entire axoneme from the base of the flagellum to its tip revealing its continuous attachment to the cell body as previously described [7]. Using higher zoom factors we can detect pro- and mature basal bodies with their associated microtubule quartet in proximity to the kDNA. The basal body is the central organizer of the cell cycle specifically involved in the duplication of the Golgi and the mitochondrion including its singular genome [8–10]. Although the precise number remains unknown, likely more than 100 proteins are involved in the biogenesis and structure of the basal body, all concentrated in a small area. Thus, precise localization of the individual components will be key for the understanding of their role in the basal body biogenesis. Centrins for example, are calcium binding proteins that are highly conserved in all centrosomes. The monoclonal antibody 20H5 that was raised against the centrin in Chlamydomonas binds to centrin 1 and 2 at the basal body and the bilobe region, respectively [8]. Applying U-ExM, we can now show that centrin is found between the maturing basal body and the associated growing microtubule quartet. To a lesser degree there is also a signal at the old basal body (Figure 2B) and at the bilobe (**\*** Figure 2B). A second very prominent candidate protein of the basal body is SAS6, a structural component of the cartwheel, that in trypanosomes is present at the pro- and the mature basal body [11] (Figure 2C). U-ExM shows that similar to the recent data from cryo-tomography SAS6 is in the center of the cartwheel and protrudes at the proximal end of the basal body [12]. Another commonly used reagent to label the basal body is the antibody YL1/2 that binds to tyrosinated α-tubulin and also cross-reacts with the basal body protein TbRP2 [13][14]. With U-ExM and at high dilutions of the primary antibody, we find tyrosinated tubulin to be more or less equally distributed at the old and new basal body. At these concentrations the apparent cross-reactivity with TbRP2 can not be seen (Figure 2D **).

### 4. Organellar proteins

One question that we liked to address was if membrane bound organelles could be analysed using U-ExM in *T. brucei*. To test this, we localized ATOM, which is part of the mitochondrial protein import machinery in the outer mitochondrial membrane, TAC102 which is a tripartite attachment complex protein inside the mitochondrion and the luminal ER component BiP [15–17]. In regular confocal microscopy ATOM is continuously distributed throughout the entire mitochondrial membrane. In expansion microscopy the distribution of the protein is visible in several hundred individual spots per cell, suggesting individual complexes can be visualized (Figure 2E). The ER resident protein BiP on the other hand seems to be more continuously distributed throughout the ER lumen even in UExM (Figure 2F), which is in good agreement with its function as a chaperone. TAC102, that was previously shown to reside inside the mitochondrion in proximity to the kDNA disc can be visualized using a monoclonal antibody (Figure 2G)

### 5. U-ExM challenges

U-ExM offers several technical advantages for visualizing ultrastructures, but it also faces significant challenges. One of the main chemicals used in U-ExM is sodium acrylate. Its quality varies from batch to batch, and impurities negatively affect polymerisation and ultimately the expansion factor. Another key challenge in expansion microscopy is the imaging process. The transparent aqueous gel starts shrinking after some time due to laser power and the evaporation of water out of the gel, which results in shifted images. Furthermore, sample storage is not as simple as for immunofluorescence slides. While immunofluorescence slides can be kept at four degrees for a long period of time, gel quality drops rapidly. It should also be noted that U-ExM requires significantly more antibody than typical immunofluorescence stainings. Also, we have experienced potentially due to the denaturation of epitopes that not all antibodies that work in regular immunofluorescence microscopy can be used in U-ExM.

## Perspective / Future directions

U-ExM enables superresolution imaging through the improvement of axial and lateral effective resolution by an isotropic increase in the distance between individual molecules to be analysed. Correspondingly, expansion by a factor of four increases resolution from around 200 nm to less than 50 nm using regular confocal microscopy and significantly further by super resolution techniques. In future, this will allow us to precisely visualize and characterize biomolecular structures that are in close proximity to each other such as components within the basal body and nuclear pore complexes. In contrast to other high resolution imaging techniques, U-ExM is accessible without the need of special equipment. In combination with protease treatment, the technique will likely allow improved intra-molecular resolution leading to in situ structure analysis, while using established reagents. Many antibodies and stains used for regular immunofluorescence microscopy show good results also in U-ExM. Furthermore, by tuning the sample functionalization steps and increasing the expansion factor, resolution in the single digit nanometer range seems feasible.

## Acknowledgements

We thank Ziyin Li and Jay Bangs for the SAS6 and BiP antibodies. André Schneider for the ATOM antibody and Keith Gull for the YL1/2 antibody. We acknowledge the Microscopy Imaging Center (MIC) of the University of Bern. The Ochsenreiter lab was supported by grants from Swiss National Science Foundation and the Uniscientia foundation.

